# Linking cells across single-cell modalities by synergistic matching of neighborhood structure

**DOI:** 10.1101/2022.04.20.488794

**Authors:** Borislav H. Hristov, Jeffrey A. Bilmes, William S. Noble

## Abstract

A wide variety of experimental methods are available to characterize different properties of single cells in a complex biosample. However, because these measurement techniques are typically destructive, researchers are often presented with complementary measurements from disjoint subsets of cells, providing a fragmented view of the cell’s biological processes. This creates a need for computational tools capable of integrating disjoint multi-omics data. Because different measurements typically do not share any features, the problem requires the integration to be done in unsupervised fashion. Recently, several methods have been proposed that project the cell measurements into a common latent space and attempt to align the corresponding low-dimensional manifolds. In this study we present an approach, Synmatch, which produces a direct matching of the cells between modalities by exploiting information about neighborhood structure in each modality. Synmatch relies on the intuition that cells which are close in one measurement space should be close in the other as well. This allows us to formulate the matching problem as a constrained supermodular optimization problem over neighborhood structures that can be solved efficiently. We show that our approach successfully matches cells in small real multi-omics datasets and performs favorably when compared to recently published state-of-the-art methods. Further, we demonstrate that Synmatch is capable of scaling to large datasets of thousands of cells. The Synmatch code and data used in this manuscript are available at https://github.com/orgs/Noble-Lab/synmatch

## 1 Introduction

Recent developments in single-cell high-throughput sequencing technologies have led to the emergence of a myriad of experimental methods that are capable of characterizing different properties of single cells in a complex biosample. For example, high-throughput sequencing methods can measure RNA expression using single-cell RNA-seq (scRNA-seq), chromatin accessibility using scATAC-seq, chromatin 3D architecture using scHi-C, and methylation profiles using scMethyl-seq. Ideally, researchers would like to be able to measure all of these properties in the same single cell in order to better understand the molecular underpinnings of the biological processes behinds cell development and disease. However, because these measurement techniques are typically destructive, frequently only complementary measurements from disjoint subsets of a given population of cells are available, providing a patchwork view of the cell’s biological processes. In such a situation, integration of the disjoint single-cell multi-omics data is critical. This has led to the rapid development of a variety of computational methods for single-cell mutli-omics integration (Adossa *et al*., 2021; Johansen and Quon, 2019; Argelaguet *et al*., 2020). What makes the problem particularly challenging however, is the fact that the different measurements, or modalities, typically do not share any features, and further, identifying correspondences between features in the domains may not be possible. Accordingly, existing methods which rely on either common cells or features across the data types cannot be applied in the fully unsupervised setting where correspondence information is absent.

This multi-modal integration problem can be generally framed in two distinct ways: (1) finding a discrete mapping between cells in the two modalities or (2) embedding the disjoint measurements into a continuous shared latent space representing the intrinsic cellular structures across cellular modalities. The generalized unsupervised manifold alignment algorithm (GUMA) (Cui *et al*., 2014), which uses a local geometry matching term, and MAGAN (Amodio and Krishnaswamy, 2018), which uses two generative adversarial networks (GANs), are examples of tools that find matchings of the cells between the two datasets. Recent methods that embed the two modalities into a common latent space and then attempt to align the embedded low-dimensional manifolds are SCOT (Demetci *et al*., 2022), Pomona (Cao *et al*., 2022a), and uniPort (Cao *et al*., 2022b) all of which employ Gromov-Wasserstein optimal transport for alignment, and MMD-MA (Singh *et al*., 2020), which aims to minimize the maximum mean discrepancy between the data sets in the latent space. Several methods rely on deep neural architectures to solve the manifold alignment task (Zuo and Chen, ????; Stark *et al*., 2020; Zhang *et al*., 2021). These methods typically use variational autoencoders as building blocks to project the data into low-dimensional manifolds and adversarial discriminators (Stark *et al*., 2020) to align the manifolds. Recently, GLUE (Cao and Gao, 2021) expanded the deep neural framework by incorporating prior knowledge about regulatory interactions to connect the feature spaces. LIGER (Welch *et al*., ????) differs from the above methods in that it employs an integrative non-negative matrix factorization approach to find the shared and dataset-specific factors across datasets in the embedded space. Finally, the UnionCom algorithm (Cao *et al*., 2020) solves both problems: it first finds a matching between distance matrices from the two modalities and then uses that matching to induce an embedding. Table 1 summarizes the recent methods based on some of their key properties. For a good review, see (Stanojevic *et al*., 2022).

**Table 1:**
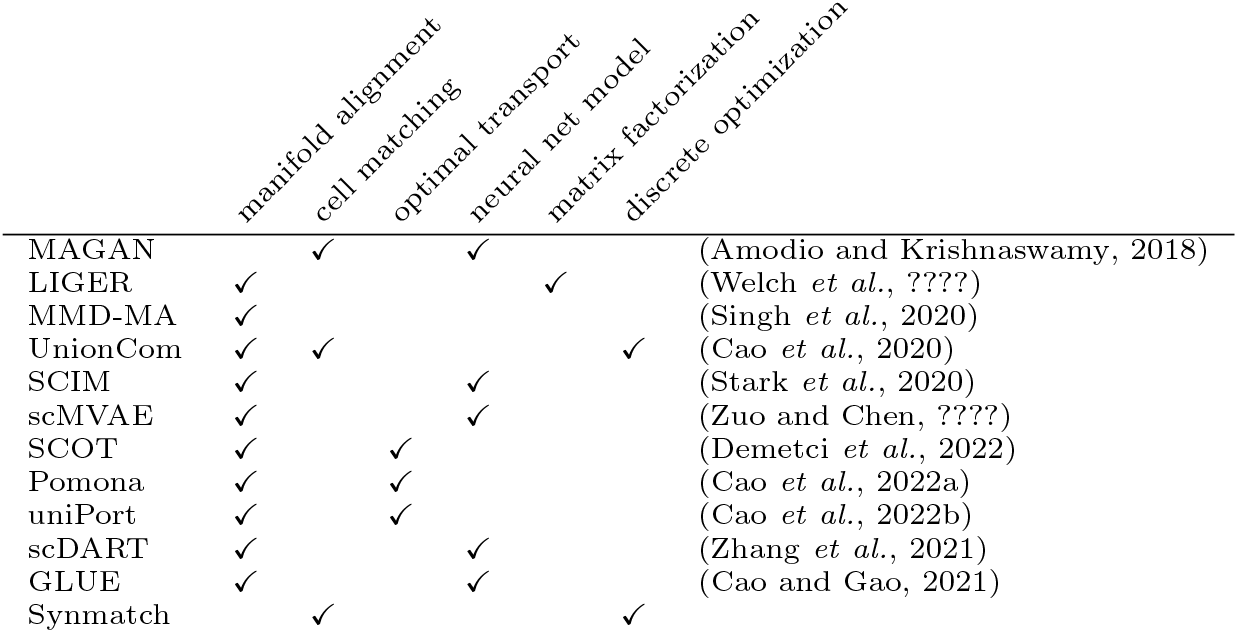
Methods for unsupervised multi-model data integration.

Here, we present Synmatch, a discrete optimization algorithm that exploits neighborhood structure and uses supermodular optimization to find a matching of the cells from two different multi-omics datasets that do not have any features in common. The key idea behind Synmatch is that the same cell, when measured in two different modalities, is likely to have similar sets of neighboring cells in the two spaces. We use this intuition to formulate the matching problem as a supermodular optimization over the neighborhood structure of the two modalities, and we solve the problem using a fast greedy heuristic that offers good theoretical guarantees. We demonstrate that Synmatch offers excellent performance in finding matchings of cells in several small single-cell multi-omics datasets, outperforming several state-of-the-art methods. We also propose an iterative procedure to allow our algorithm to scale up to datasets of thousands of cells while maintaining its excellent performance. Our work stands out from recently developed algorithms for modality integration in that it seeks direct mapping between the cells based in their shared combinatorial properties in respectively their own latent space rather than to find a common latent space within which affinity may be defined. Although Synmatch was designed to integrate single-cell multi-omics data, it is applicable to problems in other areas where matching of observations from different modalities is needed, as long as the main assumption—observations that are close (and have combinatorial properties) in one modality should be close (and thus have similar combinatorial properties) in the other modality—holds.

## 2 Methods

### 2.1 The Synmatch algorithm

Synmatch takes as input two matrices of single-cell profiles measuring different cellular properties, such as gene expression and chromatin accessibility, and outputs a matching of the cells across the datasets. Figure 1 illustrates the key concept behind Synmatch. Two similar cells that are close in one modality likely share the same biological (and thus combinatorial) properties.^1^ Hence, it is likely that they are close in the other modality, reflecting the similarity of their biological properties.

**Figure 1:**
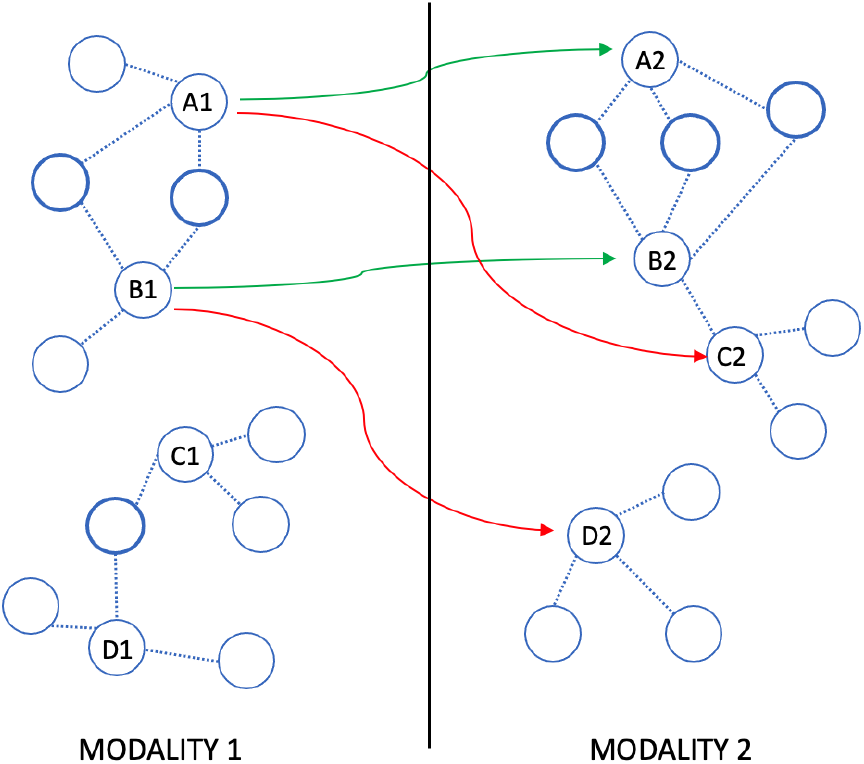
Synergistic matching of neighborhood structure. Synmatch aims to match cells that share common neighbors in each data modality. In the figure, each labeled cell is connected to its three nearest neighbors by dotted edges. The two cells A1 and B1, which have neighbors in common (indicated by thicker circles) in modality 1, should be matched (green arrows) to the two cells A2 and B2, which also share neighbors in modality 2. Conversely, A1 and B1 should not be matched (red arrows) to cells C2 and D2, which do not share neighbors. In the first step of the algorithm these common neighbors help diffusion propagate between A1—B1 and A2—B2. This in turn facilitates the optimization, which operates cooperatively on pairs of edges and aims to match pairs of cells with shared local structure across the modalities.

We denote the two sets of measurements as 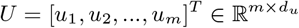 and 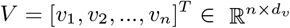, where 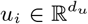 and *v_j_* ∈ *R^dv^* are column vectors describing, respectively, cell *i* and cell *j*. Our goal is to find a matching between these two sets of cells, where we describe a matching as a set *E*’ of edges in a bipartite graph between *U* and *V*: *E*′ ⊆ *E* = *U* × *V*. In the resulting matching, we require that no node in *U* or *V* has incident edge degree greater than 1. Hence, if *m* ≠ *n* then some cells in one of the two sets will be left unmatched. The Synmatch algorithm proceeds in two phases.

In the first phase, we compute a diffusion-based similarity between the cells in *U* and, separately, among the cells in *V* (so there is no direct similarity computed at this stage between any *u* ∈ *U* and *v* ∈ *V*). This measure captures both the local and global relationships among the cells in the each modality. For the moment we discuss only *U*. We use the cosine distance between any two cells *u_i_* and *u_j_* in *U* to assign a weight to the edge *e_U_* (*i*, *j*) in the complete graph *G_u_* = (*U*, *U* × *U*) induced by the cells in *U*. We choose cosine similarity instead of Euclidean because it has been shown to be a considerably more robust measure of cell-to-cell similarity (Korsunsky *et al*., 2019). Next, we employ a diffusion kernel (Kondor and Lafferty, 2002) to spread activation across the graph *G_U_*. Briefly, the Laplacian of a graph *G_u_* shifted by *λ* is defined as *L_U_* = *D_U_* + *λI* – *A_U_*, where *I* is the identity matrix, *D_U_* is the diagonal matrix *d_ii_* = ∑*_j_ e_U_* (*i*, *j*), *A_U_* is the adjacency matrix of the graph, and *λ* is a parameter controlling how far the activation spreads across the graph *G_U_*. As shown in (Qi *et al*., 2008), the amount of activation at equilibrium can be efficiently computed as 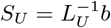, where *b* is the elementary unit vector with 1 for the nodes introducing the flow and 0 for the rest. We note that this diffusion kernel has been successfully utilized in variety of computational problems ranging from protein function prediction (Tsuda and Noble, 2004) to cancer gene identification (Hristov *et al*., 2020). We use *S_U_*(*u_i_*, *u_j_*) as a measure of the **similarity** between cells *u_i_* and *u_j_*. We analogously compute *S_V_* for the cells in *V*.

In the second phase, we construct a mapping between the cells in *U* and *V* based on *S_U_* and *S_V_*. We consider all pairs of cells (*u_i_*, *u_j_*) ∈ *U* and all pairs of cells (*v_l_*, *v_k_*) ∈ *V*. Intuitively, if cells *u_i_* and *u_j_* are close to one another in *U*, then the corresponding cells in *V* should also be close to one another. That is, a good matching is one in which a large *S_U_*(*u_i_*, *u_j_*) implies a large *S_V_*(*v_l_*, *v_k_*) and vice versa, a property well expressed by the square root of the product, i.e., 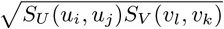. A large value of this product thereby provides evidence not only that the cells (*u_i_*, *v_l_*) should be matched but also that the cells (*u_j_*, *v_k_*) should be matched. The second part of our objective, in fact, expresses a form of complementarity between matched edges. Any given matched pairs of cells, in the form of an edge, say (*i*, *j*) ∈ *E*′, should offer benefit to all other cell pairs (*j*, *k*) ∈ *E*′ commensurate with the tendency of the corresponding cells (*j*, *k*) to be close whenever (*i*, *j*) is close. This property is expressed precisely using an objective 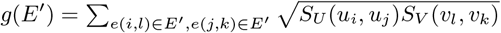 that judges the quality of the set of edges E’ being considered in a match. We note that *g*(*E*′) is a set function objective that scores any *E*′ ⊆ *E* and in fact is a well known supermodular objective (Bilmes, 2022). Of course not all subsets *E*′ ⊆ *E* are valid matchings, so this leads us to a constrained optimization problem of the form:

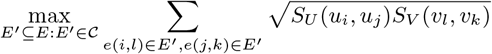

where 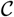 is the constraint that the edges *E*′ must form a bipartite matching. Since the function being optimized is supermodular, and as long as the diagonal is not zero, we can efficiently maximize it using a greedy heuristic which iteratively adds to *E*′ the edge that improves the objective function the most while maintaining the bipartite constraints. While supermodular maximization subject to matroid constraints is normally hard, this algorithm has a theoretical approximation guarantee (Bai and Bilmes, 2018) depending on the diagonal component of the implicitly expressed |*E*| × |*E*| matrix and depending on the curvature of the supermodular function.

### 2.2 Scaling Synmatch to large numbers of cells

Because our approach needs to examine all *O*(*m*^2^*n*^2^) possible pairs of edges, it does not immediately scale to thousands of cells due to memory constraints. In practice, Synmatch can easily run on a personal computer if |*U*| ≤ 300 and |*V*| ≤ 300 or |*U*||*V*| ≤ 10000. For larger datasets, we aggregate the cells in each modality into a small number *c* < 100 of clusters, which we refer to as a “meta-cells,” using equal size *k*-means clustering. Then, we compute pairwise similarity matrix *S_MU_* between the meta-cells in modality *U*. Specifically, for two meta-cells *m*_1_ and *m*_2_ in *U*, *S_MU_*(*m*_1_, *m*_2_) = ∑_*u_i_*∈*m*_1_, *u_j_*∈*m*_2__ *S_U_*(*u_i_*, *u_j_*)/|*m*_1_||*m*_2_|. We analogously compute *S*_MV_. We match the meta-cells using the Synmatch algorithm as described above (using *S_MU_* and *S_MV_* instead of *S_U_* and *S_V_*), and then recursively match individual cells within each pair of matched meta-cells. If a given pair of matched meta-cells contains more than a total of 10000 cells and hence cannot be matched directly, then we repeat the procedure of aggregating these cells into sub-meta-cells that we proceed to match, and so on.

### 2.3 Datasets

We use real single-cell multi-omics datasets in our analysis. All datasets are generated by co-assays; hence, we know the correct cell-to-cell correspondence for benchmarking.

The first dataset comes from the SNARE-seq assay (Chen *et al*., 2019) (accession number GSE126074) and consists of a mixture of human cell lines (BJ, H1, K562 and GM12878). The gene expression information is stored in a cell × gene counts matrix with dimensionality 1047 × 18,666 while the chromatin accessibility information is stored in a Boolean cell × peak matrix of size 1047 × 136,771. We reduce the dimensionality of the datasets in the same way as in (Singh *et al*., 2020): we apply PCA to the gene expression data and select the top 10 components, resulting in a 1047 × 10 matrix. We reduce the sparsity and noise of chromatin accessibility data by using the cisTopic (González-Blas *et al*., 2018) framework, resulting in a 1047 × 19 matrix.

The second dataset is generated by the scGEM assay (Cheow *et al*., 2016) (accession SRP077853) and simultaneously profiles gene expression and DNA methylation of human somatic cells undergoing conversion to induced pluripotent stem cells. This dataset consists of 177 cells and has dimensions 177 × 34 for the gene expression data and 177 × 27 for the chromatin accessibility data.

The third dataset is derived from the recently developed SHARE-seq assay (Ma *et al*., 2020) (accession GSE140203). It jointly profiles chromatin accessibility and gene expression in 34,774 mouse skin cells. The unprocessed data matrices have dimensionally 34,774 × 164,105 and 34,774 × 20,085, respectively. We reduce each data matrix using PCA to two matrices of sizes 34,774 × 10 each.

### 2.4 Evaluation metrics

To assess the performance of each algorithm we employ three evaluation metrics.

The FOSCTTM score is the fraction of samples closer than the true match (Liu *et al*., 2019). For a given cell *c* in one modality, we identify its correct match *m*(*c*) in the other modality. We then calculate the Euclidean distance between all of the cells in the other modality to *m*(*c*), and we compute the fraction of them that are closer to *m*(*c*) than the predicted match *p*(*c*). The final score is the average this fraction across all data points in both domains. Lower scores are better, with a score of 0 reflecting a perfect matching.

The neighborhood overlap score quantifies the percentage of all cells whose correct match lies within a given size neighborhood of the cell they have been matched to (Stanley III *et al*., 2020). Specifically, if a cell *c* is matched to cell *p*(*c*), then a neighborhood of fixed size *k* = 0, 1, 2,…, *n* around *p*(*c*) is examined whether to ascertain whether it contain the correct match *m*(*c*). For each *k*, the average of all cells from each modality for which this condition is true is reported. The score ranges from 0 to 100%, with a higher percentage being indicative of a better recovery of the cell-to-cell relationship between the two datasets.

Unlike the previous two scores, the third score, label transfer accuracy, does not require knowing the correspondence between cells in the two domains. Instead, label transfer accuracy makes use of cell type label information. This score aims to assess the ability to correctly transfer cell labels from one domain to another based on the predicted matching. As in (Cao *et al*., 2020), we train a *k*-nearest neighbor classifier (with *k* = 5) on one of the modalities, and we use it predict the cell labels in the other modality. The label transfer accuracy is the percentage of cells with correctly predicted labels. This score ranges form 0 to 100%, with a higher percentage being indicative of better performance.

### 2.5 Hyperparameter tuning

In our analysis we compare Synmatch with three state-of-the-art single-cell alignment methods, none of which uses any correspondence information: SCOT (Demetci *et al*., 2022), UnionCom (Cao *et al*., 2020), and MMD-MA (Singh *et al*., 2020). We run each competing method a over grid of its hyperparameters (trying to keep the grids about the same size of 120 points) selecting the hyperparameters that yield the lowest average FOSCTTM score. SCOT has two hyperparameters: regularization weight *ϵ* ∈ {0.0001, 0.0005, 0.001, 0.005, 0.01, 0.05, 0.1} and number of neighbors *k* ∈ {10, 20, 30, 40, 60, 80, 100, 200, 500, 1000}; MMD-MA three: weights *λ*_1_, *λ*_2_ ∈ {10^-3^, 10^-4^, 10^-5^, 10^-6^, 10^-7^} and dimensionally *p* ∈ {4, 5, 6,16, 32}; UnionCom four: trade-off *β* ∈ {0.1, 1, 10, 20}, regularization coefficient *ρ* ∈ {0, 5, 10, 15, 20}, dimensionally *p* ∈ {4, 6, 16, 32}, and *k_max_* ∈ {40, 100}. We note that the performance of these methods greatly depends on the choice of parameters, and the ones provided by default achieve significantly worse performance than optimal.

## 3 Results

### 3.1 Synmatch improves cell matching on small single-cell multi-omics datasets

First, we investigate performance of our method when it constructs a matching between cells in small datasets, when no clustering into meta-cells is necessary. We run Synmatch on scGEM co-assay data, which profiles gene expression and DNA methylation in 177 human somatic cells. This dataset was previously used to showcase the performance of the UnionCom algorithm (Cao *et al*., 2020). For comparison, we also run three state-of-the-art methods—SCOT, UnionCom, and MMDMA—on the same dataset. To judge performance we employ three metrics: (1) the neighborhood overlap (Cao *et al*., 2020), which is defined as the percentage of cells that can find their corresponding cells from the other dataset in a neighborhood of a given size around the cells that the algorithm matches them to, (2) label transfer accuracy (Johansen and Quon, 2019), which measures how well cell type labels are transferred from one dataset to another, and (3) the FOSCTTM score, which quantifies the fraction of samples closer than the true match (Singh *et al*., 2020) (see Section 2.4 for details). Hyperparameters for all methods were selected by minimizing the FOSCTTM score over a predefined grid (Section 2.5).

Our results show that Synmatch outperforms all three methods in finding the correct matching between cells across modalities. Synmatch achieves the best (i.e., lowest) FOSCTTM score of 0.19 compared to 0.20 for SCOT, 0.22 for MMD-MA and 0.23 for UnionCom. Synmatch also performs well according to the neighborhood overlap score, where it exhibits a higher score than the competing algorithms across neighborhoods of size < 50 (Figure 2A). For larger neighborhoods, all four methods perform similarly. Finally, the label transfer accuracy for all four methods is similar, with a slight edge for Synmatch: Synmatch correctly transferred 60% of cell type labels from one modality to another compared to 58%, 59%, 59% for SCOT, UnionCom, and MMD-MA, respectively.

**Figure 2:**
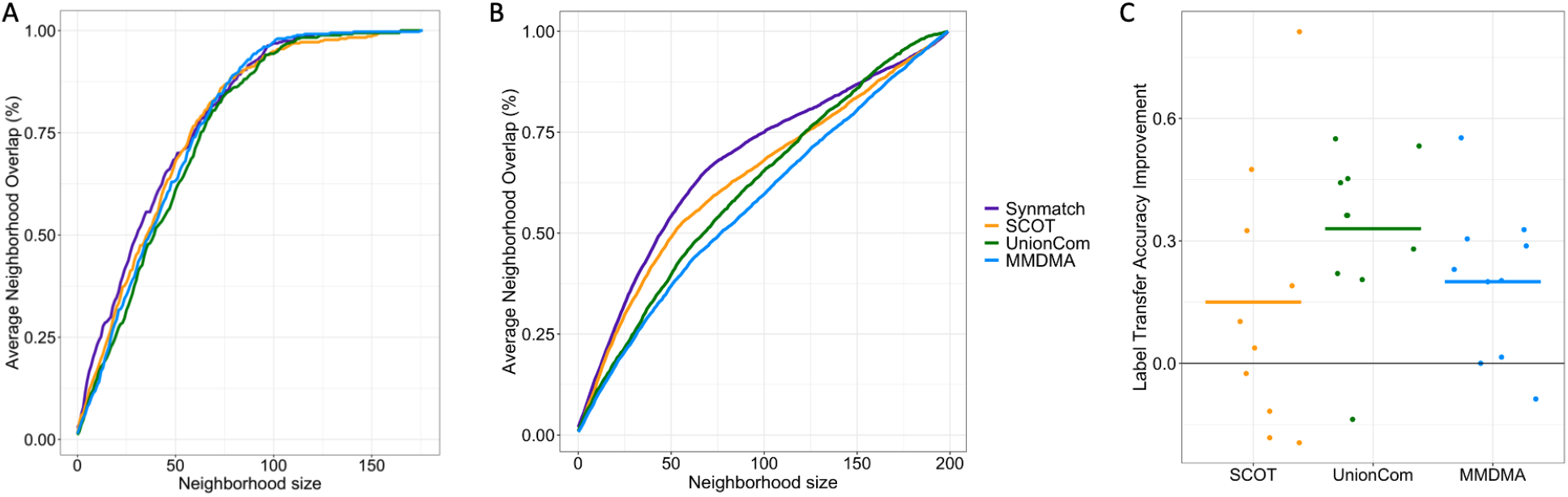
Performance comparison on small datasets. (A) The figure plots the neighborhood overlap as a function of neighborhood size on the scGEM dataset. The four series correspond to Synmatch and three other state-of-the-art methods. (B) The figure plots the neighborhood overlap, averaged over 10 different data sets of size 200, drawn from the SNARE-seq dataset, as a function of neighborhood size. (C) The figure plots, for each of three competing methods, the difference in label transfer accuracy compared to Synmatch, with positive values representing an improvement by Synmatch. Each dot corresponds to a different randomly sampled subset of size 200 from the SNARE-seq assay.

Next, we ran Synmatch on subsets of cells from a SNARE-seq co-assay dataset (Chen *et al*., 2019), which measures gene expression and chromatin accessibility. To assure robustness, we repeatedly subsampled ten dataset of size 200 cells, and we report the average performance for each algorithm. As before, we select hyperparameters by grid search, optimizing the FOSCTTM score. The neighborhood overlap for Synmatch is higher than those of the competing methods (Figure 2B). Furthermore, Synmatch excels at correctly transferring cell type labels, improving over UnionCom on average by 0.33, over MMD-MA by 0.23, and over SCOT by 0.20 (Figure 2C).

### 3.2 Synmatch successfully scales to thousands of cells

In practice, many multi-modal single-cell datasets contain thousands of cells and hence cannot be directly analyzed by Synmatch due to its memory requirements. Accordingly, we implemented and tested a recursive variant of Synmatch, which involves clustering the cells into a small number of meta-cells, matching those meta-cells with Synmatch, and then matching the cells within each pair of matched meta-cells, again with Synmatch (Section 2.2). To validate the approach, we ran Synmatch on 10 random samples of 10,000 cells drawn from a SHARE-seq co-assay (Ma *et al*., 2020), which profiles chromatin accessibility and gene expression. For the clustering step, we employed equal size K-means with *k* = 250 to group the cells into *c* = 40 meta-cells. As before, we compared Synmatch’s performance to that of SCOT, MMD-MA, and UnionCom, and we used the same hyperparameter grid search procedure.

Synmatch performs well in this comparison. Synmatch’s FOSCTTM score is the best 0.34 (0.36 for SCOT, 0.39 for MMDMA and 0.42 for UnionCom). In terms of neighborhood overlap, Synmatch is often the best-performing method (8 of 10 neighborhood sizes that we considered), and when it is not the top-ranked method it is always second-ranked (Figure 3A). Synmatch also achieves the highest label

**Figure 3:**
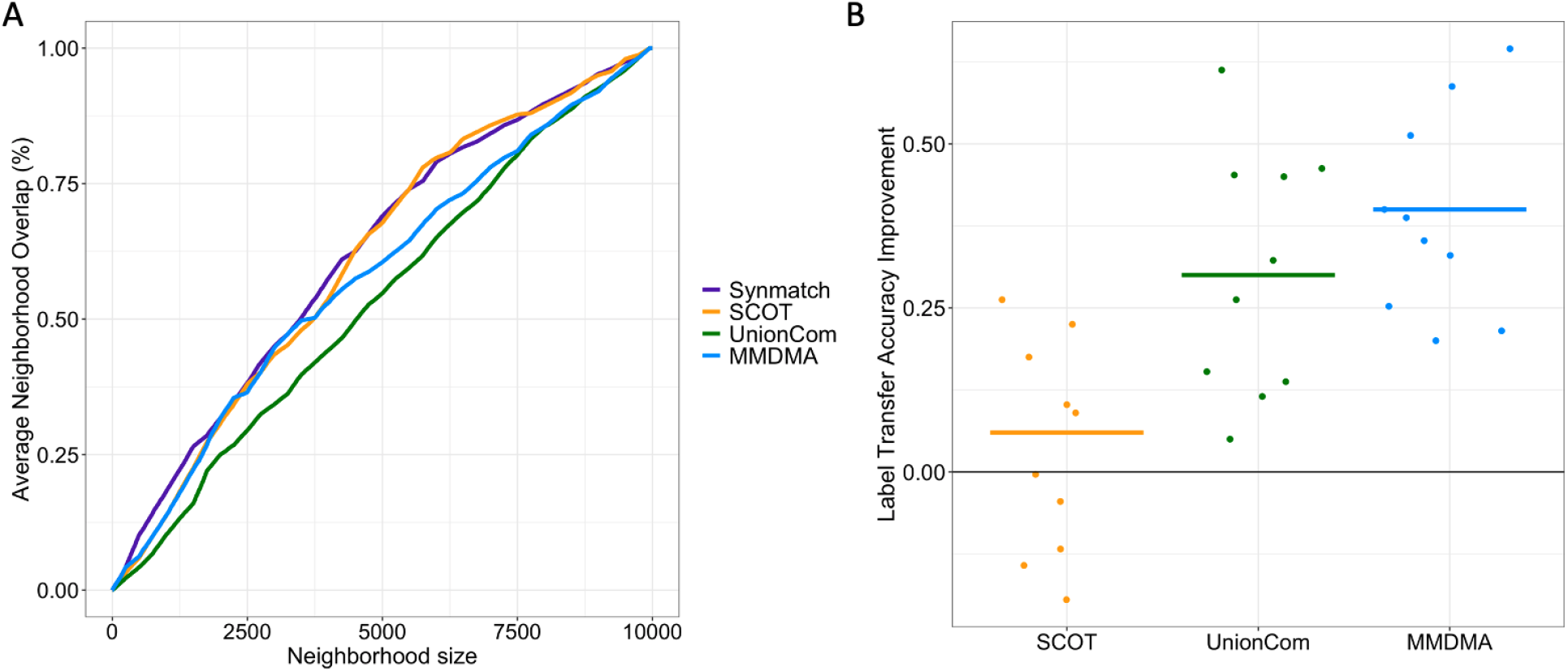
Performance comparison on large dataset. (A) The figure plots the neighborhood overlap, averaged over 10 different data sets of size 10000, drawn from the SHARE-seq dataset, as a function of neighborhood size. (B) The figure plots, for each of three competing methods, the difference in label transfer accuracy compared to Synmatch, with positive values represnting an improvement by Synmatch. Each dot corresponds to a different randomly sampled subset of size 10,000 from the SNARE-seq assay.

transfer accuracy, exceeding the second-ranked method (SCOT) by 0.06 on average. Notably, Synmatch does a better job transferring cell type labels than MMD-MA and UnionCom in all 10 runs.

### 3.3 Investigating variants of the Synmatch algorithm

There are two critical components in the process of scaling Synmatch up to larger datasets: grouping the cells into meta-cells and matching the meta-cells between modalities. Accordingly, we explore these two steps in detail. We find that both of these steps have a significant impact on the performance of Synmatch, and variations in either step can lead to very different results.

First, we test Synmatch using several other clustering strategies: regular K-means, agglomerative hierarchical clustering, and spectral clustering. We also tested SeaCells (Persad *et al*., 2022), a recently published method specifically designed for deriving meta-cells from single-cell data. We find that, on average, each of these clustering algorithms performs worse than equal-size *k*-means, when evaluated based on the FOSCTTM score (Figure 4A). Further investigation shows that these clustering methods yield very imbalanced clusters, with some clusters containing only a handful of cells and others containing hundreds. Thus, if in the second step of our algorithm a meta-cell *m*_1_ with less than 10 cells is matched to a meta-cell *m*_2_ with more than 100 cells, then the subsequent matching of the individual cells from these two meta-cells will leave the majority of the cells unmatched. We attempted to resolve this problem by returning all the unmatched cells into a common pool and re-running Synmatch on them. This approach, however, leads to both a significantly slower performance and worse overall matching.

**Figure 4:**
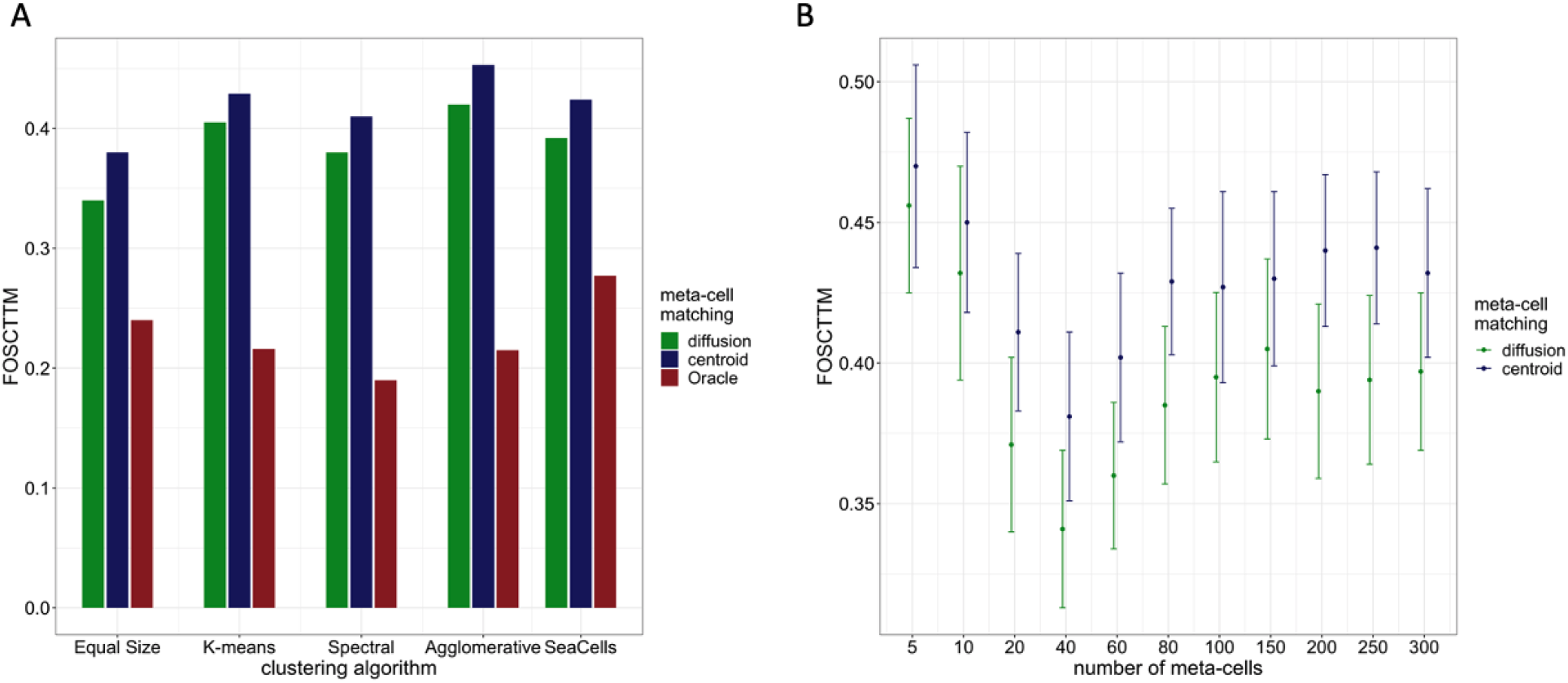
Performance comparison between different clustering and meta-cell matching strategies. (A) The figure plots the FOSCTTM score, averaged over 10 different data sets of size 10000 drawn from the SHARE-seq dataset, for several clustering approaches (groups of bars) with number of clusters *c* = 40 and three different meta-cell matching strategies (the color bars). (B) The figure plots the average FOSCTTM score over 10 different data sets of size 10000 for various numbers of meta-cells (clusters) using Equal size K-means with the two corresponding meta-cell matching strategies.

Second, we explore two different strategies to match the meta-cells to one another. The first strategy represents each meta-cell by its centroid and runs Synmatch to match the centroids. The second strategy computes an average diffusion-based similarity (*S_MU_* and *S_MV_*) between the meta-cells in each modality, which Synmatch then uses (Section 2.2). As an upper bound for comparison, we also include a third strategy: an oracle provides the true 1—1 cell correspondence to find the best possible match between the meta-cells. Briefly, for every pair of possible meta-cell matchings, the oracle computes the number of cells that can possibly find their correct match if two meta-cells are matched, uses it to assign weight on the edge between the two meta-cells, and finally uses the Edmonds Karp algorithm to find the maximum bipartite matching between the meta-cells. The average diffusion-based strategy consistently outperforms centroid-based one across different clustering strategies, including regular *k*-means (Figure 4A, blue bars are always taller than green bars) We hypothesize that the reason is that the centroids represent a crude and imperfect center of each meta-cell in each measurement space, whereas the *S_MU_* and *S_MV_* matrices better capture the similarity relationships among the meta-cells. The large gap between the oracle-based and the Synmatch-based matching strategies indicates that our method could achieve significantly better performance if the meta-cells were linked more accurately.

Third, we explore the number of meta-cells c we cluster the cells into. Because of memory constraints we require *c* ≤ 300 and test the performance of Synmatch for ten different values of *c* (Figure 4B). We observe that *c* = 40 yields the lowest average FOSCTTM score and that performance plateaus for *c* > 100. We also note that the diffusion-based linking strategy is consistently better than the centroid one for all c. Interestingly, using small number of meta-cells (*c* = 5) leads to the worst score. We suspect that the reasons for that is twofold. Given only a handful of cells Synmatch cannot leverage neighborhood structure information since there are only a few neighbors possible. Further, these meta-cells are very large in size and if they are incorrectly matched this has a major negative downstream impact on the ability to correctly match the individual cells within them.

Finally, we investigate the impact of the diffusion decay parameter *λ*. By default we use *λ* = 0.5, which balances the importance of the local and global neighborhood structures. We observe that small changes in the value of this parameter (*λ* ∈ (0.3, 0.8)) do not affect significantly Synmatch’s performance (Figure 5). However, if *λ* is set to extreme values such as 10^-4^ or 10^4^ the performance drops dramatically. This is expected since in both cases the diffusion does not capture the neighborhood structure—in the former it spreads almost uniformly across the cells and in the latter it is centered around a single cell.

**Figure 5:**
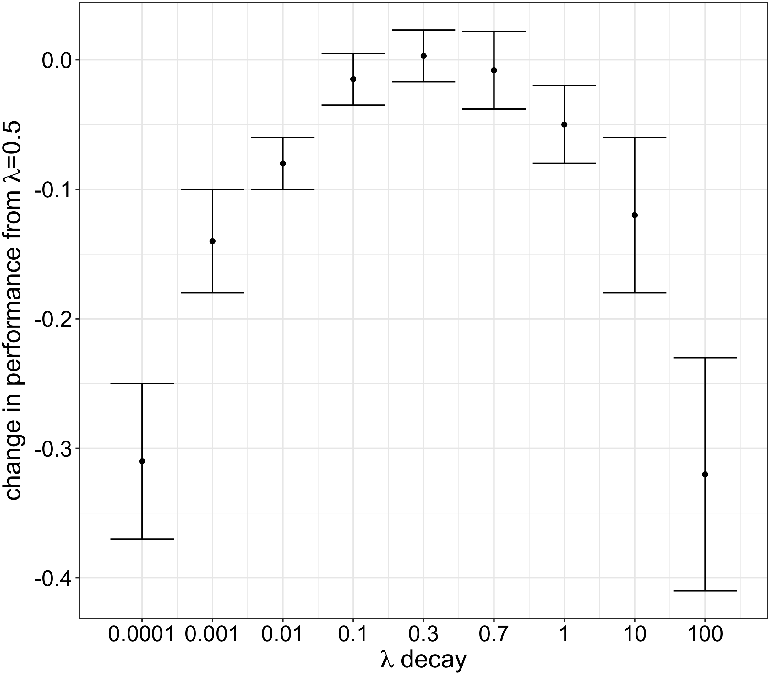
Performance depends on adequately capturing neighborhood structure. The figure plots the change in performance, averaged over 10 different data sets of size 10000, drawn from the SHARE-seq dataset, as the diffusion parameter *λ* is varied. The baseline is the default *λ* = 0.5.

## 4 Discussion

In this study we present Synmatch, an algorithm that directly maps cells across single-cell modalities that do not share any features or have any known cell-to-cell correspondence information. Synmatch exploits the neighborhood structure around the cells in each modality, seeking a matching that maps nearby cells in one modality to nearby cells in the other modality. The problem is framed as a discrete supermodular optimization and is solved efficiently. We demonstrate that Synmatch successfully matches cells in several small real single-cell multi-omics datasets and show that it can scale to large dataset of thousands of cells. Synmatch compares favorably to state-of-the-art integration methods based on three commonly employed evaluation metrics.

From a theoretical perspective, our algorithm stands out from the majority of recently published work for two reasons: (1) it finds matching of the cells directly without the need to project the two modalities into a shared latent space and (2) it uses a discrete optimization instead of the commonly employed optimal transport or deep learning auto-encoder-based architecture. As new tools for integration of single-cell omics data continue to emerge, those that aggregate cells into “super-cells” or “meta-cells” (Persad *et al*., 2022) reflecting underlying biological properties could provide a better stepping stone for scaling up our approach.

Future work should focus on finding ways to improve the matching of the meta-cells, as our results indicate that this step has a significant impact on the overall performance. One element of our approach is that it does not immediately provide soft cell mappings, e.g., when a cell in one modality is probabilistically matched to cells in the other modality. It can, however, be extended to the probabilistic case by using log-supermodular probability distributions or approaches where we exclude certain cells from a matching to arrive at score sensitivities that could be interpreted as probabilities. Our method can easily provide a many-to-one matching by relaxing the bipartite constraint to more general intersection of matroid constraints.

## Footnotes

1 In this work, “combinatorial properties” includes things such as neighborhood structure in graphs, but could, in general, include any graph property such as triadic closure, neighborhood reciprocity, and so on (Newman, 2018)

